# Using composite phenotypes to reveal hidden physiological heterogeneity and model oxygen saturation variation of high altitude acclimatization in a Chinese Han longitudinal cohort

**DOI:** 10.1101/336446

**Authors:** Yi Li, Yi Wang, Menghan Zhang, Yanyun Ma, Kun Wang, Xiaoyu Liu, Meng Hao, Chang Sun, Xiaofeng Wang, Xingdong Chen, Yao Zhang, Wenyuan Duan, Longli Kang, Bin Qiao, Jiucun Wang, Li Jin

## Abstract

Altitude acclimatization is the physiological process of the human body adjusting to the decreased availability of oxygen. Since several physiological processes are involved and the relation among them is complicated, analyses of single-traits is insufficient in revealing the complex mechanism of high altitude acclimatization. In this study, we examined whether these physiological responses could be studied as composite phenotypes which are represented by a linear combination of physiological traits. We developed a strategy which combines both spectral clustering and Partial Least Squares Path Modeling (PLSPM) to define composite phenotypes based on a cohort study of 883 Chinese Han males. And we captured 14 composite phenotypes from 28 physiological traits of high altitude acclimatization. Using these composite phenotypes, we applied k-means clustering to reveal hidden population physiological heterogeneity in high altitude acclimatization. Furthermore, we employed multivariate linear regression to systematically model (Model 1 and Model 2) oxygen saturation (SpO_2_) changes in high altitude acclimatization and evaluated the model fitness performance. And composite phenotypes based Model 2 has better fitness than single-traits based Model 1 in all measurement indices. Therefore, this new strategy of defining and applying composite phenotypes can be considered as a general strategy of complex traits research, which may also shed light on genetic loci discovery and phenome analyses.

## INTRODUCTION

Altitude acclimatization is the physiological process of the human body adjusting to the decreased availability of oxygen^1^. It comprises of several physiological responses in the body, including ventilation function, cardiac function, oxygen delivery function, hematology, muscle structure and metabolism, oxygen consumption and so on^2,3^. The most important physiological responses are in the cardiorespiratory and the hematology system^2^. Oxygen saturation (SpO_2_) reflect the most straightforward physiological changes^2,4–7^. The SpO_2_ quickly decreased in three days in lowlanders ascending directly to 4,300 m, followed by a rise over weeks at altitude^1,2,8,9^. Another well-known physiological change is the hemoglobin concentration in the blood^1,8–10^. It is also known that individuals vary in both the speed and extent to altitude acclimatization^1,11,12^. The variations of responses across individuals provide an opportunity to explore the mechanism of altitude acclimatization^1,9,11^.

Since several physiological processes are involved and the relation among them is complicated, analyses of single-traits are insufficient to capture the complex mechanism of high altitude acclimatization^1,4,9^. Therefore, analysis of composite phenotypes, i.e. combinations of physiological phenotypes could become a promising alternative^13–15^. There are several methods to extract composite phenotypes from multiple traits, such as Principal Component Analysis (PCA)-based methods^14,16,17^ and Partial Least Squares (PLS)-based methods^9,18,19^. PLS-based methods have better performance than PCA-based methods^18,19^. Partial Least Squares Path Modeling (PLSPM) is the PLS-based approach to Structural Equation Modeling^20–22^, which can also be viewed as a method for analyzing multiple relationships between groups of variables. In the PLSPM framework, there are generally two ways to define composite phenotypes, i.e., latent variables^9,19–22^: one is using the prior knowledge and the other is using data-driven methods such as spectral clustering^23,24^.

Here, we conducted a two-phase longitudinal study of high altitude acclimatization (baseline and chronic phase) in a large sample of 883 Chinese Han young males. Overall 28 physiological phenotypes were collected from these individuals at each phase. First, we extracted composite phenotypes from physiological phenotypes in high altitude acclimatization by introducing a data-driven strategy constituting spectral clustering^23,24^ and PLSPM^20,21^ algorithm. Second, using these composite phenotypes, we revealed hidden population physiological heterogeneity in high altitude acclimatization using k-means clustering^25^. Third, we modeled the changes of SpO_2_ during high altitude acclimatization using multivariate linear regression^26^, and further evaluated the advantages of composite phenotypes over single phenotypes. The workflow was summarized in **Fig. 1**, which is also the design of this study. The term phenotype used in this manuscript are referred to as “The Extended Phenotype^27^”.

**Figure 1.**
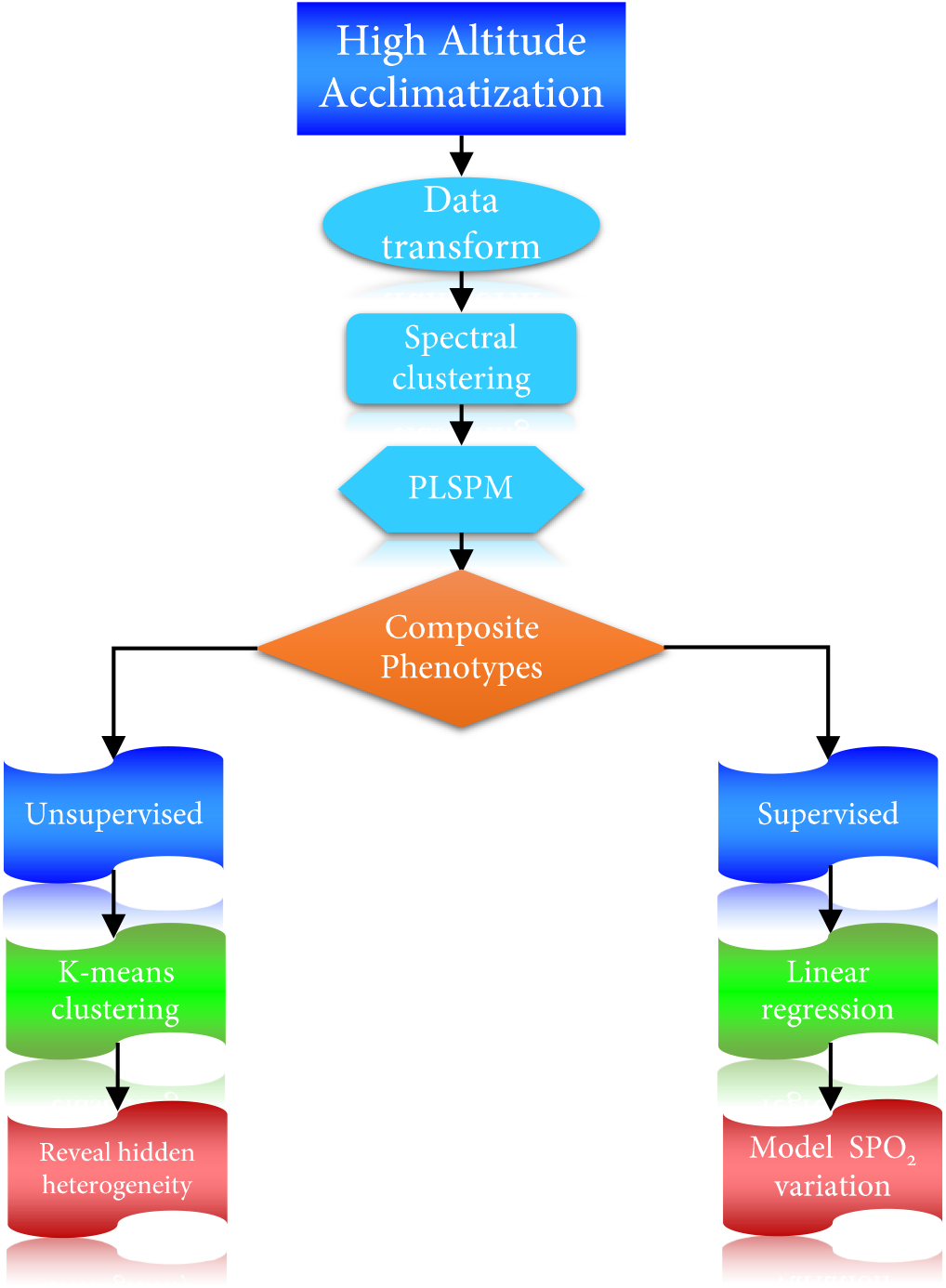
The workflow and design of this study.

## METHOD

### Study overview

To explore the physiological changes at two phases (baseline and chronic, **Table 1**) of high altitude acclimatization, the longitudinal data were transformed into change data^28^. To extract composite phenotypes from the 28 physiological traits, spectral clustering^23,24,29^ was applied firstly (**Fig. 1**). Based on the spectral clustering results (composite phenotype structure, **Fig. 2**), PLSPM^20^ was applied to construct and estimate the 14 composite phenotypes (**Table2, Fig. 3**). Using the 14 composite phenotypes, we applied k-means clustering algorithm^23,25^ to explore physiological heterogeneity in high altitude acclimatization (**Fig. 4**). To further investigate the physiological patterns of the two groups, pairwise Pearson correlation heatmap was shown (**Fig. 5**). To model how physiological traits systematically relate to SpO_2_ changes in high altitude acclimatization process, two multivariate linear regression models^26^ were constructed. Finally, to evaluate the fitness of two models, AIC^30,31^, BIC^31,32^, 10-fold CV^33^ RMSE^34^ and leave-one-out RMSE were measured (**Table 3**). In summary, firstly we have a problem in biology and then we tried to solve it using several effective statistical methods.

**Figure 2.**
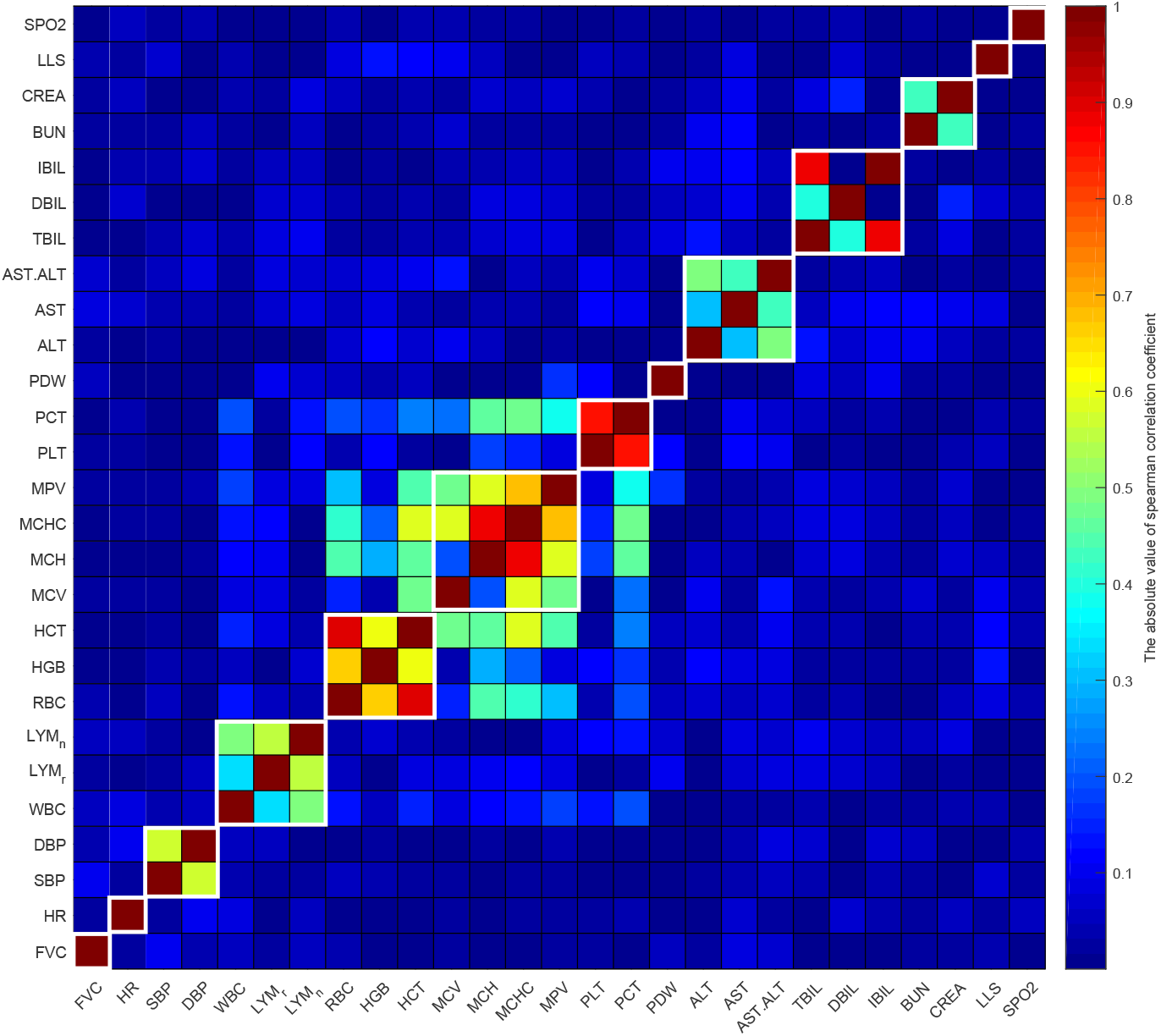
The absolute value of spearman correlation heatmap of 28 physiological phenotypes. The spearman correlation coefficient ranges from 0 (dark blue) to 1 (dark red). The spectral clustering results are marked by white boxes. For example, SBP and DBP are grouped together, and their absolute spearman correlation coefficient is about 0.6 (yellow-green color).

**Figure 3.**
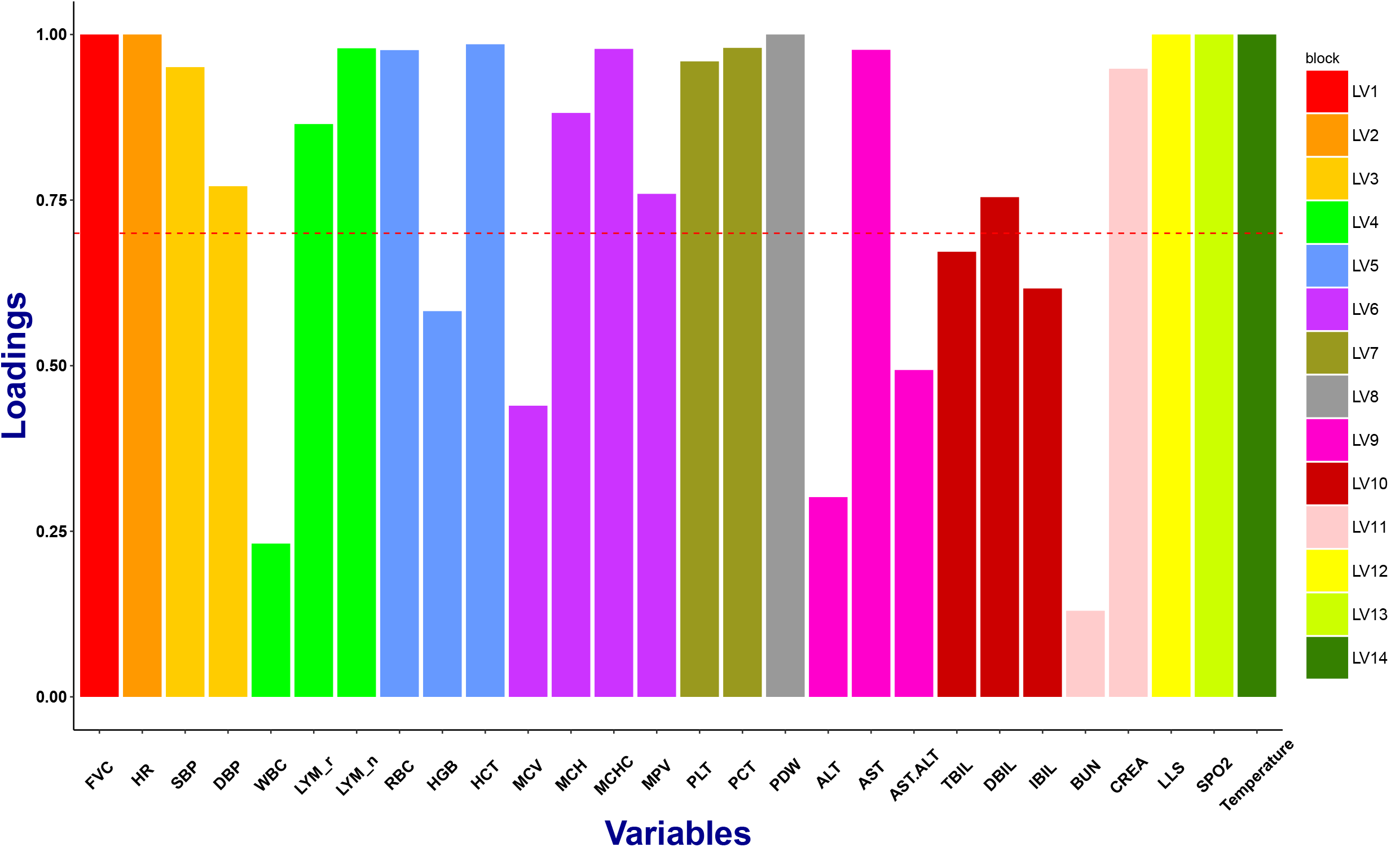
The PLSPM loadings of 14 composite phenotypes of high altitude acclimatization. The 14 composite phenotypes (LV1, …, LV14) are represented by 14 different colors, and the height of each colorful bar is the loading (correlation) of each composite phenotype. Acceptable values for the loadings are values greater than 0.7 (threshold line), indicating that more than 49% (0.7 × 0.7) of the variability in a single phenotype (like SBP or DBP) is captured by its composite phenotype (like LV3).

**Figure 4.**
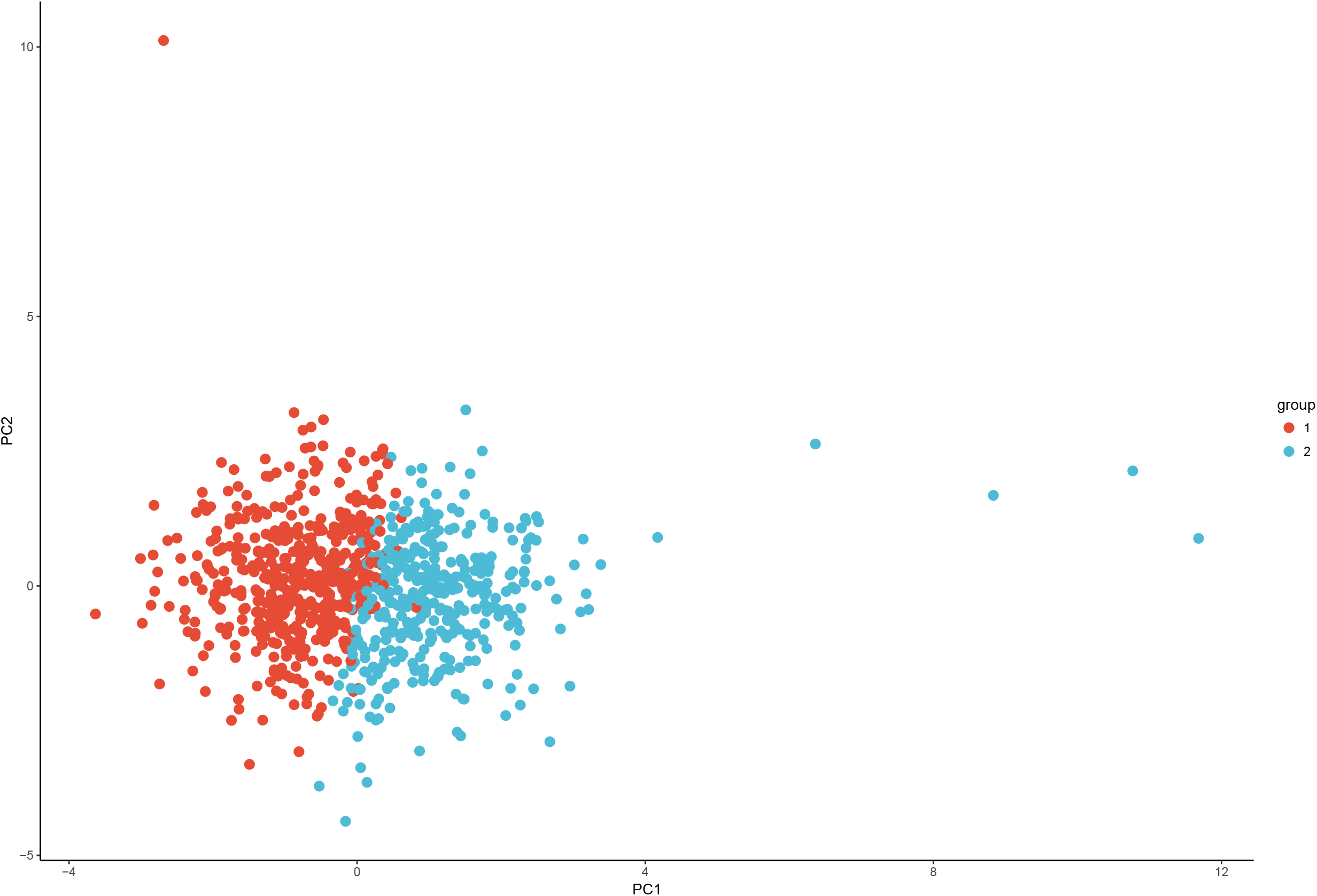
K-means clustering results on individuals using the 14 composite phenotypes (LVs). The 883 individuals are clustered as two groups (group 1 with 508 individuals and group 2 with 375 individuals) based on their composite phenotype scores. The PCA plot is just visualization of the k-means clustering results (group 1 with red color and group 2 with blue color accordingly).

**Figure 5.**
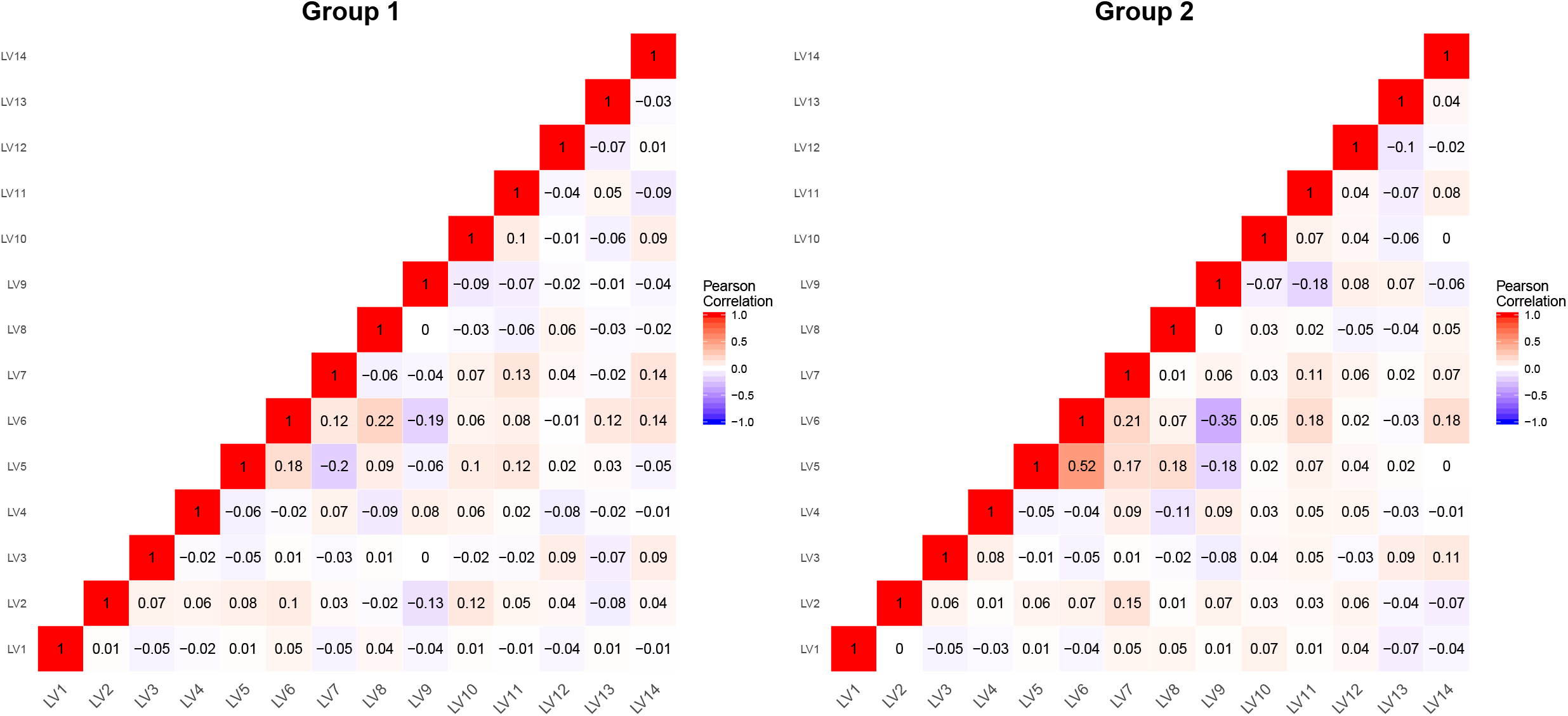
The pairwise Pearson correlation heatmap of 14 composite phenotypes (LV1,…, LV14) of two groups. The Pearson correlation coefficient ranges from −1 (blue) to 1(red). The left figure represents the Pearson correlation heatmap of 14 LVs of group 1 and the right figure represents the Pearson correlation heatmap of 14 LVs of group 2.

**Table 1.** The 28 physiological traits from 883 Chinese Han young males at baseline and chronic phases of high altitude acclimatization.

| Variables | Baseline |  | Chronic |  | Pvalue* |
| --- | --- | --- | --- | --- | --- |
|  | Mean | Sd | Mean | Sd |  |
| ALT | 19.11 | 8.14 | 15.02 | 9.08 | **6.28E-45** |
| AST | 15.64 | 5.83 | 46.05 | 15.24 | **2.33E-139** |
| AST.ALT | 0.85 | 0.18 | 3.91 | 2.54 | **2.14E-139** |
| TBIL | 11.94 | 1.77 | 14.51 | 24.90 | **1.41E-10** |
| DBIL | 2.69 | 0.61 | 5.70 | 1.70 | **3.72E-136** |
| IBIL | 9.25 | 1.24 | 8.81 | 24.95 | **1.96E-39** |
| BUN | 5.08 | 1.12 | 6.21 | 1.17 | **3.20E-87** |
| CREA | 58.32 | 10.38 | 113.02 | 12.20 | **2.85E-139** |
| WBC | 6.21 | 1.34 | 8.21 | 1.70 | **1.55E-125** |
| LYM% | 36.16 | 7.48 | 40.61 | 10.71 | **7.71E-42** |
| LYM# | 2.21 | 0.52 | 3.31 | 0.99 | **1.78E-106** |
| RBC | 4.88 | 0.39 | 5.73 | 0.48 | **5.25E-144** |
| HGB | 150.15 | 10.05 | 179.59 | 13.46 | **2.07E-144** |
| HCT | 0.44 | 0.03 | 0.50 | 0.04 | **1.35E-136** |
| MCV | 90.60 | 5.14 | 87.39 | 4.89 | **6.49E-126** |
| MCH | 30.91 | 2.34 | 31.39 | 2.18 | **1.16E-21** |
| MCHC | 341.07 | 17.94 | 359.14 | 16.24 | **3.97E-85** |
| PLT | 206.88 | 42.79 | 258.94 | 51.49 | **2.18E-124** |
| PCT | 2.01 | 0.42 | 2.72 | 0.51 | **9.95E-139** |
| MPV | 9.78 | 1.26 | 10.54 | 0.62 | **9.59E-68** |
| PDW | 13.79 | 2.25 | 18.03 | 2.53 | **9.71E-142** |
| FVC | 444.45 | 38.26 | 412.31 | 59.16 | **2.03E-51** |
| SBP | 110.88 | 10.45 | 124.34 | 12.94 | **2.51E-87** |
| DBP | 73.21 | 8.65 | 75.98 | 9.61 | **7.46E-13** |
| HR | 66.49 | 9.57 | 87.16 | 10.99 | **3.24E-123** |
| Temperature | 36.22 | 0.12 | 36.38 | 0.29 | **1.75E-39** |
| SPO2 | 97.76 | 2.08 | 85.82 | 3.80 | **6.64E-132** |
| LLS | 0.88 | 1.59 | 1.40 | 1.73 | **1.14E-10** |

\* Pvalues were calculated by Wilcoxon Rank-Sum Test (paired= true).
The significant (under Bonferroni correction) pvalues were shown in bold.

**Table 2.** PLSPM composite phenotypes unidimensionality evaluation.

|  | Biological Meanings | Manifest variables | Mode | MVs | C.alpha | DG.rho | eig.1st | eig.2nd |
| --- | --- | --- | --- | --- | --- | --- | --- | --- |
| **LV1** | forced vital capacity | FVC | A | 1 | 1.00 | 1.00 | 1.00 | 0.00 |
| **LV2** | heart rate | HR | A | 1 | 1.00 | 1.00 | 1.00 | 0.00 |
| **LV3** | blood pressure | SBP, DBP | A | 2 | 0.70 | 0.87 | 1.54 | 0.46 |
| **LV4** | immune system | LYM#, LYM%, WBC | A | 3 | 0.55 | 0.77 | 1.76 | 1.21 |
| **LV5** | number of red cells | RBC, HCT, HGB | A | 3 | 0.89 | 0.93 | 2.45 | 0.48 |
| **LV6** | hemoglobin concentration | MCH, MCHC, MPV, MCV | A | 4 | 0.79 | 0.87 | 2.52 | 1.01 |
| **LV7** | number of platelets | PLT, PCT | A | 2 | 0.94 | 0.97 | 1.88 | 0.12 |
| **LV8** | platelet distribution width | PDW | A | 1 | 1.00 | 1.00 | 1.00 | 0.00 |
| **LV9** | liver function | ALT, AST, AST/ALT | A | 3 | 0.25 | 0.04 | 1.35 | 1.31 |
| **LV10** | bilirubin | TBIL, DBIL, IBIL | A | 3 | 0.59 | 0.79 | 2.00 | 1.00 |
| **LV11** | renal function | BUN, CREA | A | 2 | 0.61 | 0.84 | 1.44 | 0.56 |
| **LV12** | Lake Louise score | LLS | A | 1 | 1.00 | 1.00 | 1.00 | 0.00 |
| **LV13** | oxygen saturation | SPO<sub>2</sub> | A | 1 | 1.00 | 1.00 | 1.00 | 0.00 |
| **LV14** | body temperature | body temperature | A | 1 | 1.00 | 1.00 | 1.00 | 0.00 |

Note: Overall 14 composite phenotypes are shown as the latent variables (**LV1, LV2...LV13, LV14**).

**Table 3.** Evaluation the goodness of fit of two multivariate linear regression models.

|  | Model1 (original variables) | Model2 (LV) | Pvalue* |
| --- | --- | --- | --- |
| AIC | 5089 | **5069** | — |
| BIC | 5227 | **5141** | — |
| 10 fold CV RMSE | 4.32 | **4.26** | — |
| 10 fold CV MSE (SD) | 18.64 (5.60) | **18.12 (5.71)** | **0.00488** |
| Leave one out CV RMSE | 4.32 | **4.265** | — |
| Leave one out CV MSE (SD) | 18.66 (53.42) | **18.19 (53.32)** | **0.00294** |

\*The Pvalues were calculated by Wilcoxon Rank-Sum Test.
The measurement indices with better fitness were shown in bold.
The significant pvalues (pvalue<0.05) were marked as bold and red color.

All the computation process of this study were realized in R (v3.3.1)^35^ and the related figures were generated by Matlab (R2015b)^36^, ‘ggplot2’^37^, ‘igraph’^38^ R packages. The computation process of composite phenotype scores was completed by ‘plspm’^20^ R package. The k-means clustering was completed by ‘factoextra’^39^ and ‘NbClust’^40^ R package. The multivariate linear regression models were calculated by ‘stats’ R package.

### Exploring relationship of phenotypes by spectral clustering

The longitudinal data of high altitude acclimatization were firstly transformed into change data^28^. All the 28 physiological traits have significant (under Bonferroni correction^41^) changes from baseline to chronic phase at 4,300m highland. And the p values were calculated by Wilcoxon Rank-Sum Test^42^ (**Table 1**). Based on the change data of high altitude acclimatization, spectral clustering^23,24^ was applied. The similarity matrixs in this study were the absolute values of spearman correlation coefficient^43^ matrixs of 28 physiological changes from baseline to chronic phase for high altitude acclimatization. The affinity matrixs were computed by applying a k-nearest neighbor filter^44^ to build a representation of a graph connecting just the closest dataset points. To compute the graph Laplacian matrix, there is also a need to get the degree matrix where each diagonal value is the degree of the respective vertex and all other positions are zero^24^. To choose the best number of spectral clustering, the eigenvalue gap (difference between consecutive eigenvalues of Laplacian matrix, **Supplementary Fig. 1**) was maximized^29^. And the spectral clustering results were the composite phenotype structure (**Fig.1** and **Fig. 2**).

### Defining composite phenotypes by PLSPM

Based on the composite phenotype structure, PLSPM^20,21^ was further applied to construct composite phenotypes. And the latent variable scores^20,22^ were calculated to estimate these composite phenotypes. As the 28 physiological traits were clustered as 14 groups, there were also 14 composite phenotypes (LV1, LV2… LV13, LV14) accordingly. PLSPM is claimed to explain at best the residual variance of the latent variables and potentially also of the manifest variables in any regression run in the model without strong assumptions^22^. To check the PLSPM blocks’ unidimensionality, the Cronbach’s alpha, Dillon-Goldstein’s rho and the first eigenvalue of the indicators’ correlation matrix were calculated^20,22^. Each composite phenotype captures a specific aspect of high altitude acclimatization (**Table 2, Fig. 3** and **Supplementary Fig. 2**).

### Revealing physiological heterogeneity by k-means

Based on the 14 composite phenotypes, k-means clustering was applied to explore physiological heterogeneity in high altitude acclimatization (**Fig. 4**). The optimal number of clusters is 2 (**Supplementary Fig. 3**) following the majority rule of 26 indices^40^. The silhouette plot (**Supplementary Fig. 4**) for k-means clustering also showed that observations are well clustered^45^. Thus the 883 Chinese Han young males were clustered into two groups (group1 with 508 individuals and group2 with 375 individuals, **Fig. 4**) based on the 14 composite phenotypes of high altitude acclimatization. To further investigate the physiological patterns of the two groups, pairwise Pearson correlation^46^ heatmap was shown (**Fig. 5**).

### Modeling oxygen saturation variation by multivariate linear regression

To model how physiological traits systematically relate to SpO_2_ changes in high altitude acclimatization process, two multivariate linear regression models^26^ were constructed. Model 1 is constructed by original 28 physiological traits changes from baseline to chronic phase at 4300m highland and SpO_2_ is the dependent variable (Y). Model 2 is constructed by 13 composite phenotypes (excluding LV13, i.e., SpO_2_) of high altitude acclimatization, and SpO_2_ is still the dependent variable (Y). To evaluate the fitness of two models, AIC^30,31^, BIC^31,32^, 10-fold CV^33^ RMSE^34^ and leave-one-out RMSE were measured (**Table 3**). We also employed Wilcoxon Rank-Sum Test^42^ to compare the 10-fold CV MSE and leave-one-out MSE of two models (Model 1 and Model 2).

### Participants

We conducted a longitudinal cohort measurement design to investigate the responses of 28 physiological traits during high altitude acclimatization. The studied subjects were first assembled at a location with an altitude of 50 m (in China) for 10–14 days, and then they arrived at highland of above 4,300 m (in China) by train. The study is comprised of two phases: baseline phase (before going to highland) and chronic phase (living at highland for about 1 month). A structured questionnaire and physiological examination for the subjects were carried out at two phases of high altitude acclimatization respectively. The subjects with cancer, diabetes and coronary heart disease were not included in this study. Overall 883 healthy Chinese Han young males aged from 17 to 36 years old were recruited. The research was approved by the Human Ethics Committee of Fudan University and written informed consent was obtained from each participant and their guardians over 18 years old.

### Physiological measurements

All the subjects (883 samples, 28 traits) were measured by physicians in General Hospital of Jinan Military Region, who were previously trained to administer a questionnaire and a physical examination. The systolic blood pressures (SBP) and diastolic blood pressures (DBP) were calculated by mean of twice measurement of a standardized mercury sphygmomanometer. Maximal vital capacity (FVC) were measured by SPIDA5. Heart rate (HR) was measured by mean of twice radial pulse, SPO_2_ was measured by Nellcor NPB-40. The body temperature (Temperature) was measured by thermometer. The blood specimens were drawn after overnight fasting for complete blood count measurement by three classification haemacytometer analyzer (Model CA-800; CIS, Japan). The blood routine indices include red blood cell count (RBC, ×10^12^/L), hemoglobin (HGB, g/L), hematocrit (HCT, %), mean corpuscular volume (MCV, fL), mean corpuscular hemoglobin (MCH, pg), mean corpuscular hemoglobin concentration (MCHC, g/L), white blood cell counts (WBC, ×10^9^/L), lymphocyte percentage (LYM%), absolute lymphocyte count (LYM#, ×10^9^/L), blood platelet (PLT, × 10^9^/L), mean platelet volume (MPV, fL), plateletcrit (PCT, fL), platelet distribution width (PDW, fL). The blood biochemical indices were measured by the automatic biochemical analyzer (Model 7060; Hitachi Ltd., Japan), including glutamate pyruvate transaminase (ALT, U/L), glutamic oxalacetic transaminase (AST, U/L), total bilirubin (TBIL, umol/L), direct bilirubin (DBIL, umol/L), blood urea nitrogen (BUN, mmol/L) and creatinine (CREA, umol/L). AST/ALT ratio and indirect bilirubin (IBIL, umol/L) were calculated indices. The Lake Louise score (LLS) system scores^47^ were also collected in two phases. The LLS questionnaire consists of five items: headache, dizziness, gastrointestinal symptoms, fatigue/weakness and difficulty sleeping. Each item was rated on a four-point scale (0= not at all, 1=mild, 2=moderate and 3=severe). Single item scores are added up and the maximal score is 15.

## RESULTS

### Exploring relationship of phenotypes in high altitude acclimatization

In this study, we collected the 28 physiological traits from 883 Chinese Han young males at baseline (before going to highland) and chronic (living at highland for about 1 month) phases of high altitude acclimatization (**Table 1**). The studied subjects were first assembled at a location with an altitude of 50 m (in China) for 10–14 days, and then they arrived at highland of above 4,300 m (in China) by train. The subjects with cancer, diabetes and coronary heart disease were not included in this study. Overall 883 healthy Chinese Han young males aged from 17 to 36 years old were recruited. All the 28 physiological traits show significant (Bonferroni correction) changes from baseline to chronic phase at 4,300m highland. These results indicate that a series of physiological phenotypes are involved in high altitude acclimatization process^1,2,9^. Since we are mainly concerned on the changes of these phenotypes, the longitudinal data were transformed into change data^28^ using Measure_chronic-baseline_ = Measure_chronic_ - Measure_baseline_. These data were used in subsequent analyses.

By analyzing the correlation between pairwise phenotypes, we found the phenotypes are structured (**Fig. 2**). For example, RBC, HGB and HCT have strong correlation with each other; and RBC almost has no correlation with LLS and SPO_2_. To further explore the relationship of phenotypes, spectral clustering algorithm^23,24^ was applied to group these 28 physiological phenotypes. To determine the number of clusters, the one with maximum eigenvalue gap (**Supplementary Fig. 1**) was chosen^29^. The correlation heatmap (**Fig. 2**) showed the spectral clustering results of 28 physiological phenotypes, and they were clustered into 14 groups (i.e. composite phenotype structure, **Fig. 2**).

### Defining composite phenotypes of high altitude acclimatization

Based on the revealed aforementioned structure, PLSPM^20–22^ was applied to extract composite phenotypes of high altitude acclimatization. Overall 14 composite phenotypes were extracted as the latent variables^20^ (LV1, LV2… LV13, LV14). Each composite phenotype is a linear combination of their manifest variables^21^, and captures a specific aspect of high altitude acclimatization (**Fig. 3, Table 2** and **Supplementary Fig. 2**). The LV5 explained the variance of RBC, HCT and HGB, which mainly reflect the number of red cells (Dillon-Goldstein’s rho = 0.93, **Table 2 and Fig. 3**). The Dillon-Goldstein’s rho focuses on the variance of the sum of variables in the block of latent variable^20,22^. The LV6 explained the variance of MCH, MCHC, MPV and MCV, which reflect the hemoglobin concentration. As the changes of MCH and MCHC were negatively related to MPV and MCV, we changed both MCH and MCHC signs to keep loadings positive^20^. The LV12 is equivalent to single-phenotype LLS and the LV13 represents single-phenotype SPO_2_.

### Revealing physiological heterogeneity in high altitude acclimatization

To explore physiological heterogeneity in high altitude acclimatization, we applied k-means clustering algorithm^23,25^ on individuals using the 14 composite phenotypes. Thus the 883 individuals could be clustered into two groups (group 1 with 508 individuals and group 2 with 375 individuals, **Fig. 4, Supplementary Fig. 4** and **Supplementary Fig. 5**) based on the 14 composite phenotypes of high altitude acclimatization. The separation of two groups of the individuals are mainly contributed by hemoglobin concentration (LV6, Wilcox Rank Sum test’s pvalue = 3.36 × 10^−90^), number of red cells (LV5) and number of platelets (LV7) (**Supplementary Table 1** and **Supplementary Fig. 5**). The results demonstrate physiological heterogeneity in high altitude acclimatization among these sampled individuals, especially in the phenotypes related with oxygen carrying capacity^1,48,49^ including hemoglobin concentration, number of red cells, platelet counts and so on. The increases in red cell number and hemoglobin concentration improve the oxygen carrying capacity of the blood to compensate for the reduction in oxygen saturation^1,50,51^.

To further characterize the relationship of the 14 composite phenotypes in each group, we calculated the pairwise Pearson correlation^46^ (**Fig. 5**). For instance, there is significant correlation (Pearson’s r = 0.12, pvalue = 0.006, **Supplementary Table 2**) between LV6 and LV13 in group 1, but no correlation between them in group 2 (Pearson’s r = −0.03, pvalue = 0.51, **Supplementary Table 3**). To compare the difference of these two Pearson correlation coefficient, fisher’s z transformation^52–55^ were applied (pvalue=0.02, **Supplementary Table 4**). There is negative correlation (Pearson’s r = −0.2) between LV5 and LV7 in group 1, while in contrast, we observed positive correlation (Pearson’s r = 0.17, fisher’s z transformation pvalue =5.58 × 10^−8^) between them in group 2. Thus we can compare the correlation networks of multiple physiological traits intuitively and focus on composite phenotypes not their manifest variables.

### Modeling oxygen saturation variation of high altitude acclimatization

Oxygen saturation (SpO_2_) reflect the most straightforward physiological change of high altitude acclimatization^2,4,5,7^. To model how other physiological traits systematically relate to SpO_2_ changes in high altitude acclimatization process, we constructed two multivariate linear regression models. Model 1 is constructed by original 28 physiological traits changes from baseline to chronic phase at 4,300m highland and SpO_2_ is the dependent variable (Y). To compare with this model, Model 2 is constructed by 13 composite phenotypes (excluding LV13, i.e., SpO_2_) of high altitude acclimatization.

To evaluate the goodness of fit of two models, the Akaike information criterion (AIC)^30,31^, Bayesian information criterion (BIC)^31,32^, 10-fold cross validation (CV)^33^ root-mean-square error (RMSE)^34^ and leave-one-out RMSE were measured (**Table 3**). Model 2 has better fitness than Model 1 in all measurement indices (**Table 3**), suggesting that the composite phenotypes is better performed in capturing variation of high altitude acclimation. From the multivariate regression result of Model 2, we also found that LV12 (LLS) is the most significant (β = −0.29, pvalue = 0.04, **Supplementary Table 5**) trait that influence SpO_2_. SpO_2_ has been well studied as predictors/indicators of AMS or LLS^1,2,4,10,56–59^. And those individuals who successfully maintain their oxygen saturation at rest, most likely do not develop AMS^2,4,57^.

## DISCUSSION

In this study, we developed a data-driven strategy (**Fig. 1**) to extract composite phenotypes from multiple physiological phenotypes of high altitude acclimatization in a large-scale Chinese Han longitudinal cohort. We first explore the relationship of phenotypes of high-altitude acclimatization. And then we extracted 14 composite phenotypes from 28 physiological traits changes of high-altitude acclimatization. This strategy could be applied to other complex traits, for example, immune diseases, cardio metabolic traits or other complex diseases.

Altitude acclimatization comprises a number of physiological responses to mitigate the effects of hypoxia^1,7^. There are many methods to analyze longitudinal data, such as linear mixed models^60^ and data transformation^28^. Since we are mainly concerned on the changes of these phenotypes, transforming the longitudinal data into change data is also a promising alternative^2,9,28,61^. Thus the transformed data was used in this study.

Since individual single-traits are insufficient to reflect the complex mechanism of high altitude acclimatization^1,4,9^, analysis of composite phenotypes could become a promising alternative^13–15^. Among several methods, PLS-based composite phenotypes have relatively interpretable biological meanings^9^. In particular, PLSPM can also be viewed as a method for analyzing multiple relationships between blocks of variables^20^.

Generally, there are two ways to define composite phenotypes in PLSPM framework^9,22^: one is using the prior specific domain knowledge and the other is using some data-driven methods like clustering. In our study, we employed the generalized standard spectral clustering^23,24^ to find the composite phenotype structure (**Fig. 2**) for high-altitude acclimatization.

This study included 28 physiological phenotypes which covered respiratory function, cardiac function, oxygen delivery function, hematology, oxygen saturation, kidney function, liver function, LLS and so on. However, there are still traits not involved in this study, such as muscle metabolism, oxygen consumption, electrocardiogram, electroencephalogram, organism metabolism and so on. And the data of this study was collected at two time points of high altitude acclimatization, which may be incomplete. The subjects in this study are all young males, the physiological responses of females may be quite different. More importantly, other factors such as genetic variations, should be studied to further understand potential physiological mechanism of high altitude acclimatization^4,7^.

In summary, we have developed a strategy constituting both spectral clustering and PLSPM to define composite phenotypes. And we effectively used this strategy to capture 14 composite phenotypes from 28 physiological phenotypes of high altitude acclimatization. The 14 composite phenotypes have clear meaning in physiology and explain most of variance in statistics. Based on these composite phenotypes, we first observed physiological heterogeneity among individuals in high altitude acclimatization. In addition, we compared the performance of composite phenotypes and regular phenotypes in predicting SpO_2_ changes. Both analyses showed that the composite phenotypes is better performed in capturing variation of high altitude acclimation. To conclude, this new strategy of defining and applying composite phenotypes can be considered as a general strategy of complex traits research^62^, especially in phenome analyses^63,64^.

## ACKNOWLEDGEMENTS

This research was supported by National Science Foundation of China (31330038, 31521003, 31460286), Ministry of Science and Technology (2015FY1117000), Science and Technology Committee of Shanghai Municipality (16JC1400500), Shanghai Municipal Science and Technology Major Project (2017SHZDZX01) and the 111 Project (B13016). The computations involved in this study were supported by the Fudan University High-End Computing Center.

## CONTRIBUTIONS

YL and LJ conceived the idea and contributed to writing of the paper. YL, YW, MHZ and LJ contributed the theoretical analysis. YL, YYM, KW, YZ, LLK, XDC, WYD, BQ, JCW and LJ contributed the data collection and data cleaning. YL, YW, MHZ, YYM, KW, XYL, WLP, CS, JCW and LJ contributed to scientific discussion and manuscript writing. YL and LJ contributed to final revision of the paper.

---

**Supplementary Figure 1.**
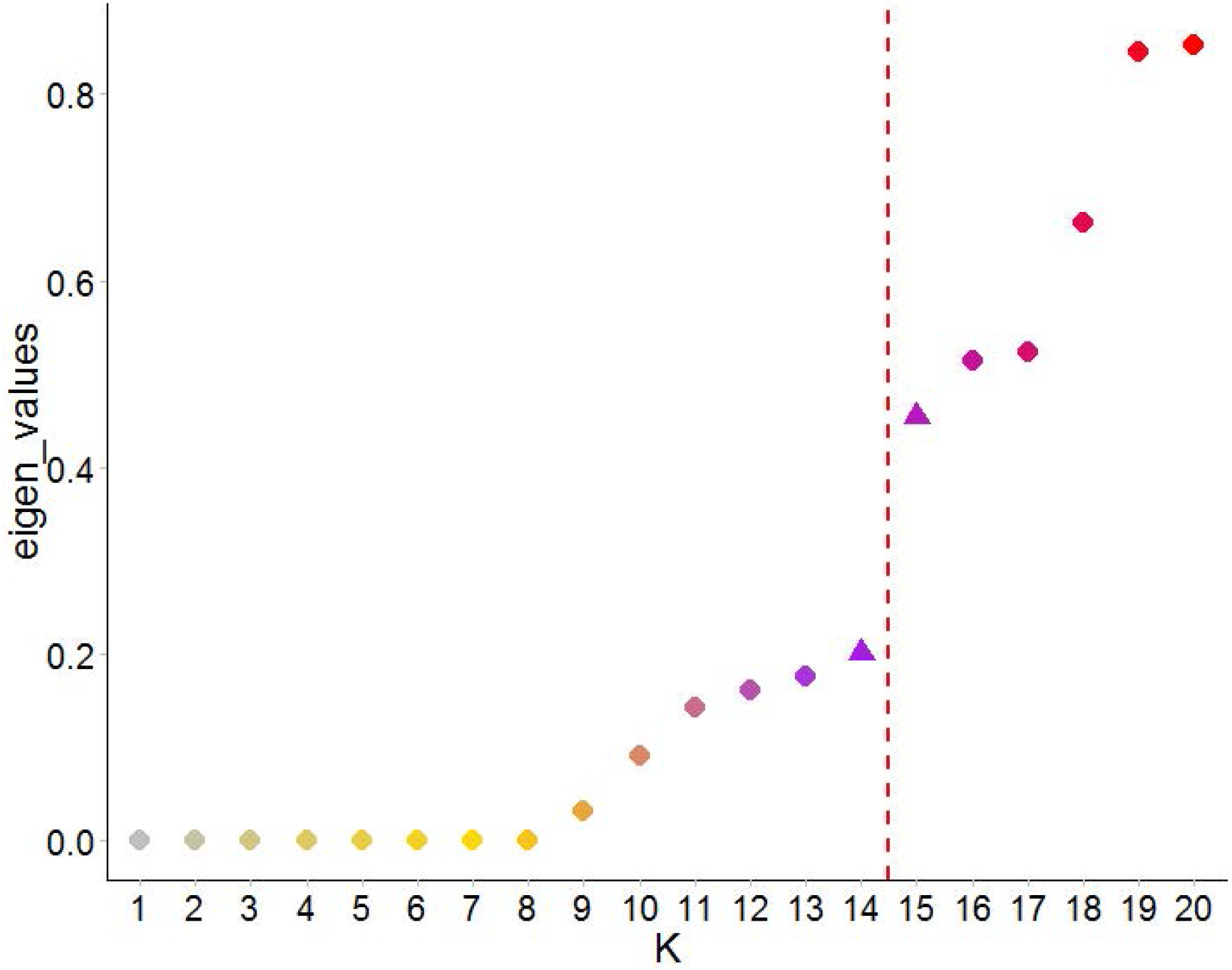
The eigenvalue gap of spectral clustering. The eigenvalue gap was maximized to choose the best number of spectral clustering (red line). And the best clustering number is 14.

**Supplementary Figure 2.**
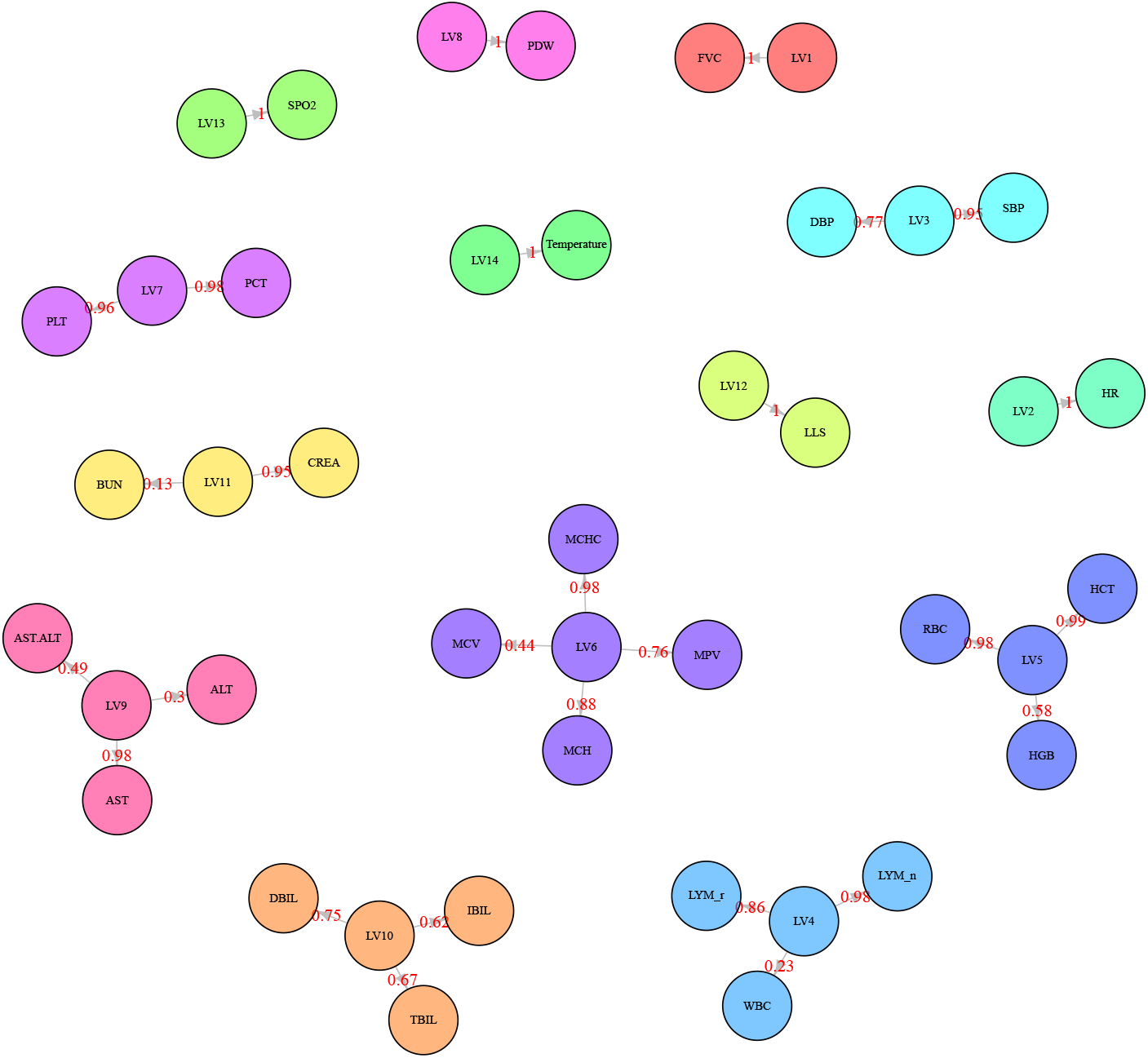
The 14 composite phenotypes of high altitude acclimatization. The numbers on the arrows are the PLSPM loadings of 14 composite phenotypes, which is the same as Figure 3.

**Supplementary Figure 3.**
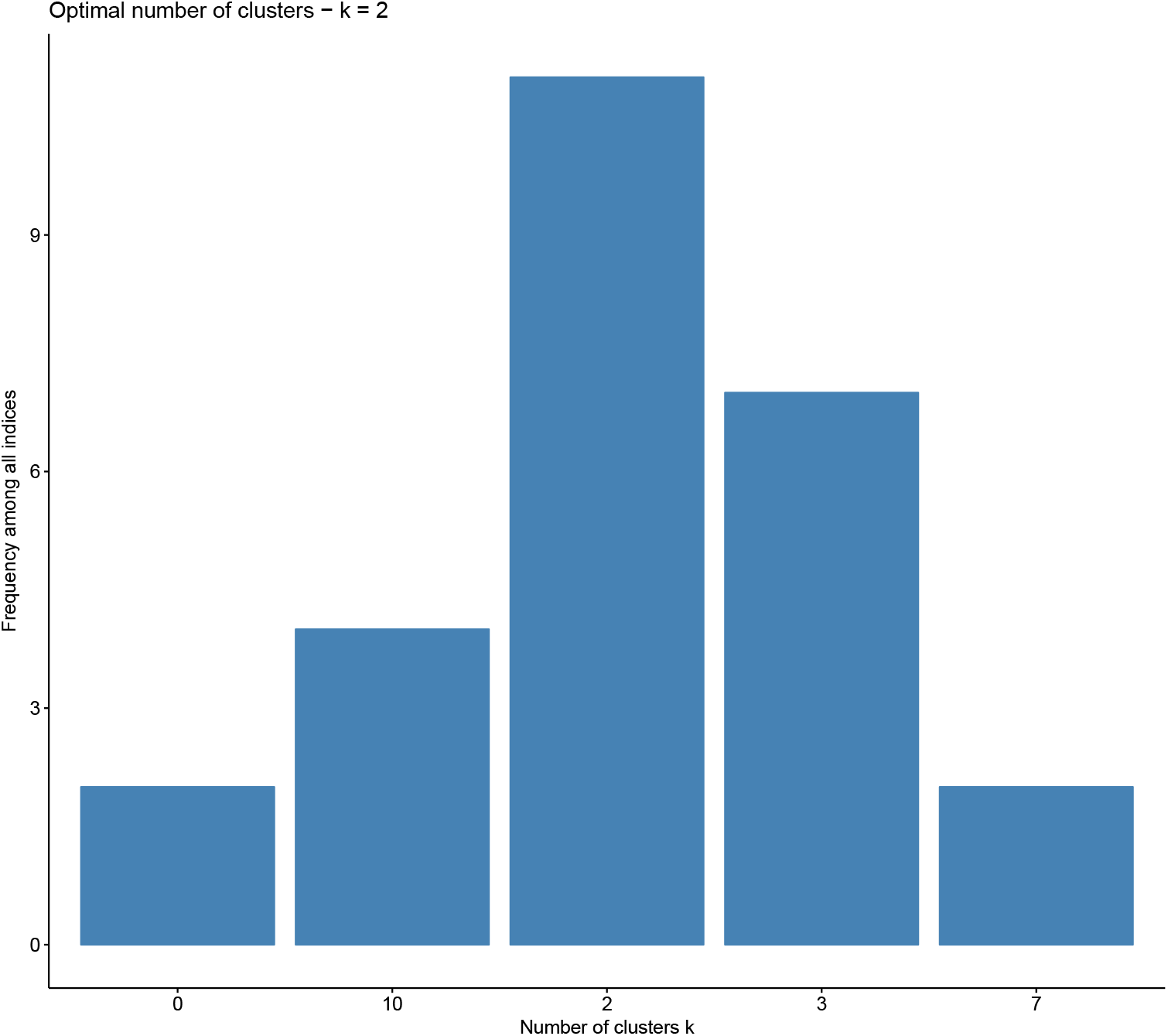
The optimal number of k-means clustering on individuals. The optimal number of clusters is 2 following the majority rule of total 26 clustering indices.

**Supplementary Figure 4.**
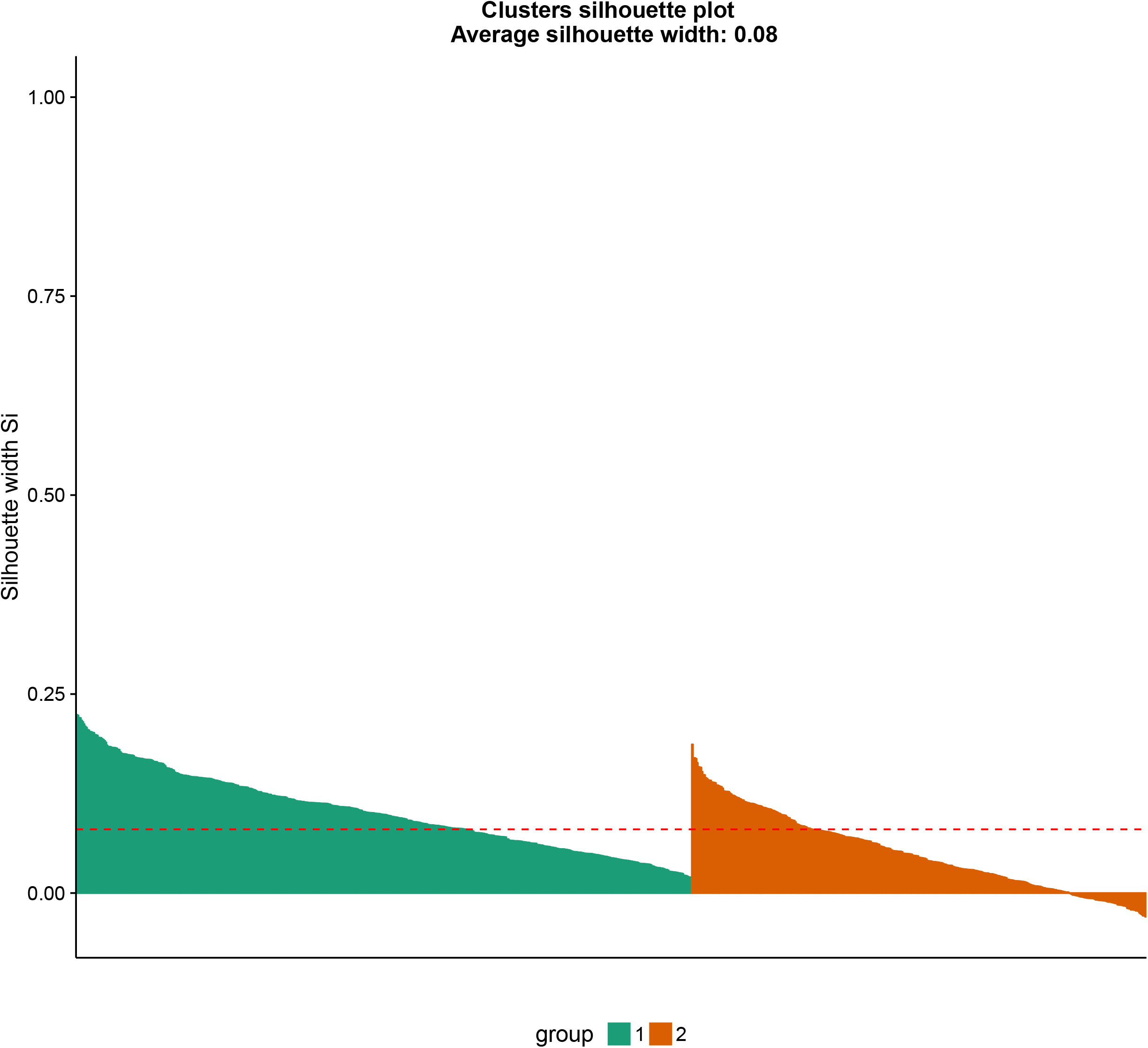
The silhouette plot of k-means clustering on individuals. Silhouette values range from 1 to −1, when silhouette value is close to 1 indicating that the individuals are well clustered. The silhouette plot for k-means clustering showed that observations are well clustered.

**Supplementary Figure 5.**
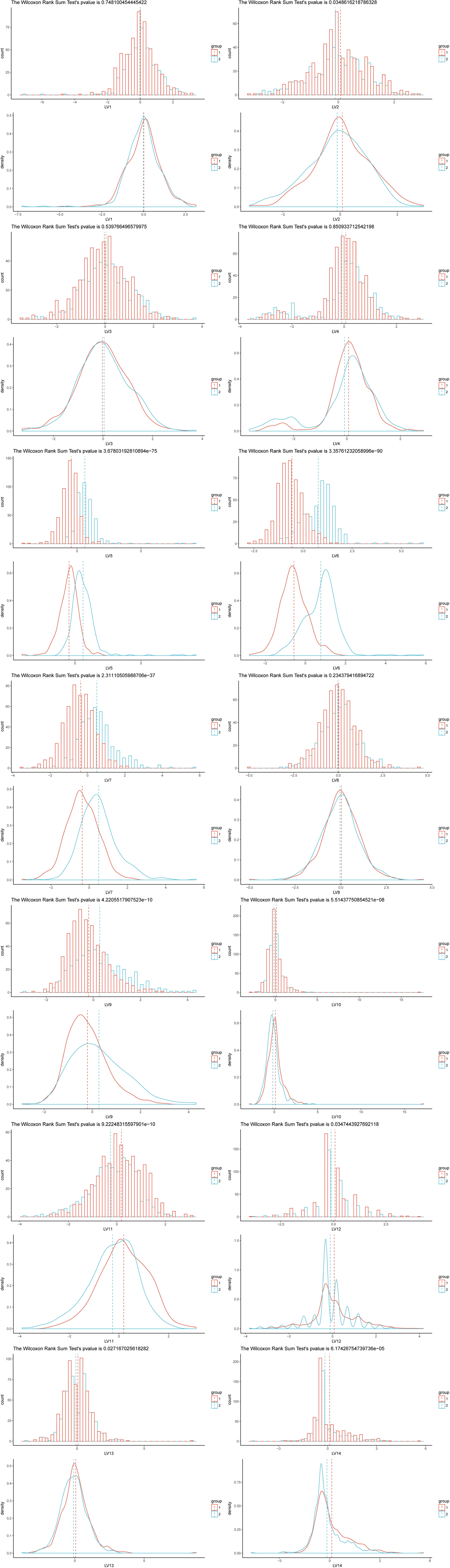
The histogram plot and density plot of each LV (14 LVs) between 2 groups (group 1 with red color and group 2 with blue color). And we further calculated the Wilcoxon Rank Sum test pvalue (the title of histogram plot) of each LV between 2 groups.

